# Nanowire-based biosensor for short DNA using fluorescent silver nanoclusters

**DOI:** 10.1101/2025.03.06.641789

**Authors:** Ivan N. Unksov, Rubina Davtyan, Christelle N. Prinz, Heiner Linke

**Affiliations:** NanoLund, Lund University, Box 118, 22100 Lund, Sweden; Solid State Physics, Lund University, Box 118, 22100 Lund, Sweden

**Keywords:** short DNA, silver nanoclusters, nanowires, fluorescence, III−V, biosensing

## Abstract

Sensitive detection of short nucleic acids is used to identify viral and bacterial diseases, detect biomarkers of cancer, as well as in gene expression studies. Currently available techniques such as PCR, electrochemical detection and SPR are typically costly and often require amplification of the DNA. Additionally, the PCR methods that involve enzymatic elongation of primers are often not optimal for short nucleic acids as a short target limits the size and specificity of the primers. Here, we demonstrate a sensing system for picomolar detection of short single-stranded DNA by fluorescence without any need for amplification, thermal cycling and expensive reagents. The platform harnesses the capability of waveguiding semiconductor nanowires to substantially enhance the signal of surface-bound fluorescent molecules. Employing molecular beacons based on DNA-templated silver nanoclusters that exhibit a larger signal in the presence of the target DNA, we improve the limit of detection by five orders of magnitude compared to flat substrates and demonstrate detection of HIV-1 DNA. The signal indicates single-molecule sensitivity of detection. Our sensor is easily adaptable for other short DNA and potentially can be mass-produced. The method requires only a small volume of analyte sample and a microscope for the detection of fluorescence on nanowires.

## Introduction

Detection of short nucleic acids at low concentrations (picomolar and below) is crucial for diagnosis of viral and bacterial diseases (Li et al., 2023), for cancer biomarker tests (Zhao et al., 2023), in gene expression studies, and for development of drugs for gene therapy (Harikai et al., 2022; Miao et al., 2024). However, the targets shorter than 40-50 nucleobases are challenging for PCR techniques. Indeed, very short primers may result in reduced specificity (Grunenwald, 2003), and their detection of short targets therefore requires additional methodological advancements (Harikai et al., 2022; Lim et al., 2021). Furthermore, there are other process- and cost-related limitations associated to PCR. It can reach attomolar limits of detection (Bruce et al., 2020; Harikai et al., 2022; Iwanaga, 2022; Nakano et al., 2017) for conventional targets but requires costly equipment, involves sample purification, enzymes, primers, amplification through thermal cycling, and fluorescently labelled DNA if optical readout is used (Madadelahi et al., 2024). As an alternative, sensors based on isothermal amplification also reach attomolar to picomolar sensitivity but remain resource demanding in terms of process, primers, and fluorophores (Kellner et al., 2019; Zhao et al., 2015). Electrochemical sensors can be used at down to femtomolar concentrations but also require highly specialized instruments and rely on electroactive labels or electrochemical changes in DNA, such as guanine oxidation, which can be non-specific (Hai et al., 2020). Surface plasmon resonance (SPR) provides label-free detection of comparable concentrations but requires costly equipment and sensor chips, while its sensitivity typically is low for short DNA targets due to their low weight (Diao et al., 2018; Li et al., 2007). Thus, alternatives are desired.

Fluorescent DNA-templated metal nanoclusters can be used in an optical biosensor based on nucleic acid hybridization. Such a biosensor enables amplification-free detection of single-stranded DNA (and potentially RNA) without synthetic fluorescent or electrochemical labels and can be applied for assessing the presence of viral and disease-associated nucleic acids in a sample. The preparation process is as simple as hybridization with a nanocluster-templating DNA, also produced in a cost-effective one-pot reaction. Self-assembling constructs based on this effect are referred to as nanocluster beacons (NCBs), pioneered by Yeh, Petty and co-workers (Obliosca et al., 2014; Yeh et al., 2012). For a NCB biosensor where fluorimetry was used for detection in bulk solution, 200 pM sensitivity was demonstrated and a limit of detection was calculated to reach tens of pM (Zou et al., 2019). However, to compete with PCR, a higher sensitivity is desired. Furthermore, a sensitive fluorimeter typically requires a relatively large sample volume of at least several hundred µL.

Here, we show that the sensitivity of a NCB sensor can be increased by using semiconductor nanowires (NWs) (Figure 1). NWs are high aspect ratio nanostructures that enhance fluorescence of surface-bound emitters due to a combination of waveguiding with directed emission (Frederiksen et al., 2017, 2016; Verardo et al., 2018), excitation enhancement (Unksov et al., 2023), and quantum yield enhancement (Sorokina et al., 2022), providing single-molecule sensitivity with a regular fluorescence microscope (Valderas-Gutiérrez et al., 2025, 2022; Verardo et al., 2019). The sensitivity can be improved by single-emitter localization, with summing the intensities of individual NWs in a field of view offering an additional improvement (Davtyan et al., 2024). The combination of DNA-templated silver nanoclusters and semiconductor nanorods, similar to NWs, has been used for the first time by Lee and co-workers for detection of ATP (Shrivastava et al., 2017). We demonstrate detection of HIV-1 DNA at picomolar concentrations and discuss practical advantages of this approach that by design can be easily adapted to other nucleic acid molecules.

**Figure 1.**
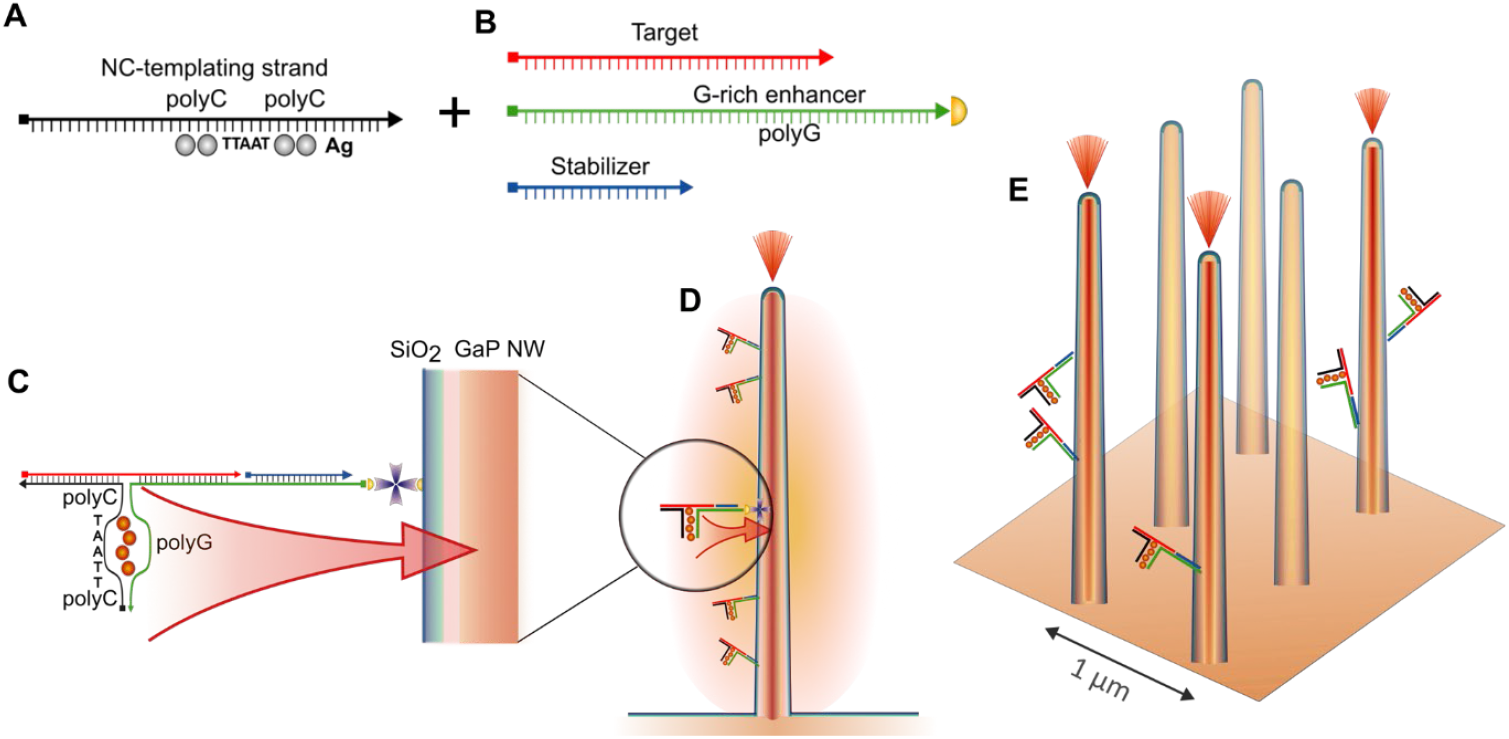
Assembly of the NCB. Nanoclusters are formed on a templating strand (A), then mixed with (B) biotinylated G-rich enhancer, target and stabilizer strands. (C) The formed NCBs are attached via streptavidin (purple cross) to a biotinylated oxide layer on a GaP nanowire. (D) Fluorescence of multiple NCBs is enhanced by the NW and guided towards the NW tip. (E) Regularly spaced NWs, a fraction of which has bound NCBs and guides the fluorescence. The distance of about 1 µm between NWs allows for imaging them as individual bright spots.

### Approach

DNA-templated metal nanoclusters are unique clusters of metal atoms that self-assemble on DNA molecules and can exhibit fluorescence depending on the DNA template sequence. Fluorescence is thus achieved without costly fluorophore modifications of the DNA, can be internalized not only at the ends but also within the macromolecule, and can be tuned by its sequence. The nanoclusters assemble in a solution containing Ag(I) ions which bind to nucleobases (Schultz et al., 2019; Swasey et al., 2015). The fluorescence of many nanoclusters is significantly increased (so called activation (Obliosca et al., 2014, 2013; Yeh et al., 2012)) when the templating strand is in proximity of a G-rich strand. This is due to the high affinity (Schultz et al., 2019; Swasey et al., 2015) of Ag(I) ions to guanine, which presumably allows for a stabilizing pocket around a cluster (Kuo et al., 2022). By varying the length, flanking nucleotides, and position of the templating strand relatively to the G-rich enhancer, a range of systems with green to infrared fluorescence have been developed (Obliosca et al., 2014, 2013; Yeh et al., 2012). In the past decade, understanding of nanocluster assembly on DNA has been improved using large-scale screening of templating oligonucleotides with subsequent template optimization by machine learning (Copp et al., 2020, 2018, 2014; Kuo et al., 2022), and X-ray crystallography for structural studies (Cerretani et al., 2019; Huard et al., 2019). The nanoclusters that were optimized in terms of their fluorescence properties are promising for biosensing.

In nanocluster beacon (NCB) biosensors, the templating and G-rich strands are by design partially complementary to a target DNA which brings them in proximity of each other (Figure 1A-C). A NCB is relatively easily assembled from regular short oligonucleotides. An advantage of NCBs is that the signal is altered or disappears in the case of even one single-nucleotide replacement in the target DNA when this replacement repositions the G-rich enhancer relatively to the templating strand (Yeh et al., 2012).

We adapted a templating sequence and a G-rich enhancer, the combination of which has recently been shown to template nanoclusters with bright red fluorescence (Kuo et al., 2022). As a target for detection, we use a 33 nucleobase DNA from the HIV-1 genome, identical to that previously tested in an electrochemical biosensor (Zhang et al., 2010), a fluorescence biosensor based on a DNA hairpin (Vet et al., 1999), and similar to the sequences detected with another nanocluster-based biosensor (Zou et al., 2019). The limit of detection in these studies was established at tens (Zou et al., 2019) and hundreds (Zhang et al., 2010) of pM.

We used gallium phosphide (GaP) epitaxial nanowires which have previously been identified as optimal for fluorescence enhancement (Unksov et al., 2023; Valderas-Gutiérrez et al., 2022). GaP is a III−V semiconductor with a high refractive index that enables waveguiding and enhancement of fluorescence (Aspnes and Studna, 1983). Due to an indirect bandgap of 2.26 eV, it also features low absorbance of visible light, minimizing signal losses and NW photoluminescence. To attach DNA to the NWs, we use a biotinylated strand that serves also as a G-rich enhancer for NCB fluorescence (Figure 1). Thus, the NCB binds to a NW only when the target brings together the templating and enhancer strands.

## Methods

### Nanowire growth and functionalization

Hexagonal arrays of GaP NWs were grown by Aligned Bio AB using MOVPE from Au seed nanoparticles deposited on a (111)B GaP substrate by displacement Talbot lithography (Coulon et al., 2019; Johansson et al., 2024; Solak et al., 2011; Wen et al., 2013). These NWs had a diameter 118 ± 5 nm (measured by SEM at half-height of the NWs) and length of 2.5 ± 0.3 µm. The average density of the NWs was 1.2 µm^-2^ (Figures 1 and 2). Each GaP NW platform was 2.5 mm × 2.5 mm × 0.3 mm in size, which was achieved by dicing. These NW platforms were subsequently coated with 10 nm of SiO_2_ using an atomic layer deposition tool (Fiji, Cambridge NanoTech). The same treatment was used for planar GaP reference platforms. The NW platforms and planar GaP coated with SiO_2_ substrates were fixed in flow channels (sticky-Slide VI 0.4, ibidi GmbH) using double-sided tape (3M). The channels were sealed with a #1.5 glass coverslip and incubated on a platform rocker for 15 minutes with biotin-bovine serum albumin (biotin-BSA, Sigma-Aldrich) at 30 µM in phosphate buffer saline (PBS) at pH 7.0. Unbound biotin-BSA was subsequently washed away with PBS, and the channels were incubated for 50 minutes with streptavidin at 1 mg/mL in PBS, which was then washed away before adding the NCBs.

**Figure 2.**
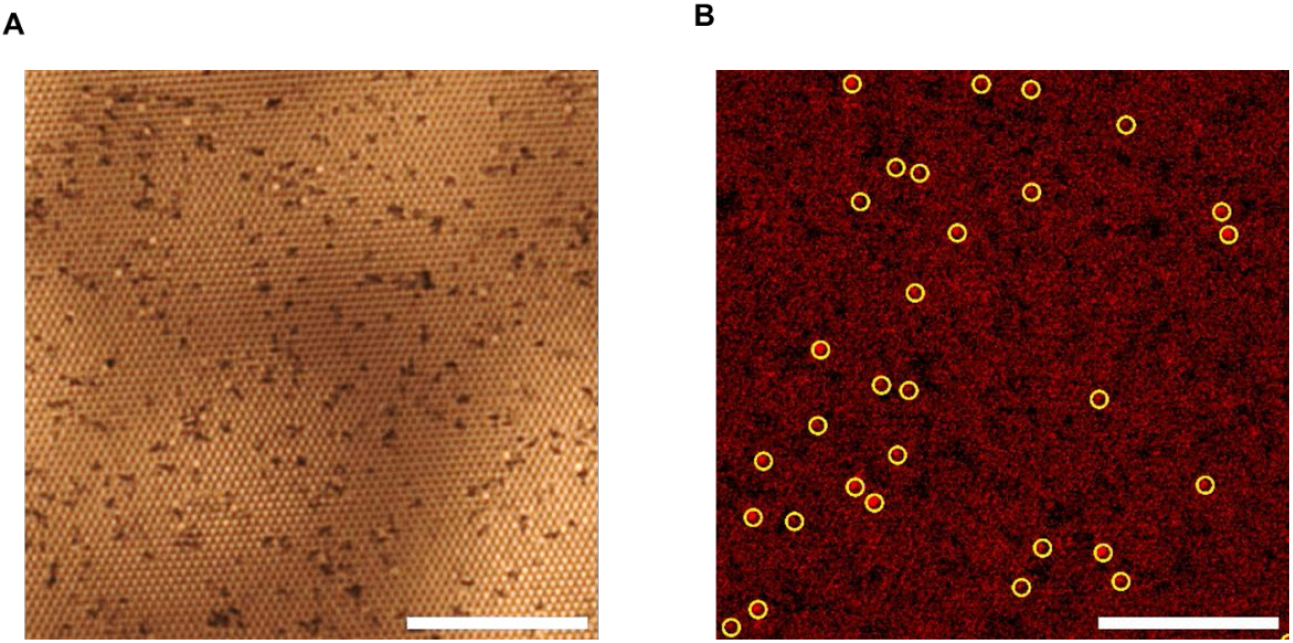
(A) NWs observed as brighter spots in a regular pattern from top view upon brightfield illumination. Dark spots are defects in the NW arrays. (B) After the NCBs have been added in presence of the target (here at 100 pM) and then unbound components were washed away, a number of NWs exhibit distinct fluorescence. Localized bright NWs are encircled in yellow, the other NWs are considered to be background by the detection procedure. For visualization only, illumination unevenness was corrected using Fourier transform, and contrast was adjusted. Scale bar is 20 µm.

### Assembling the NCBs

We used similar nanocluster-templating and G-rich enhancer DNA sequences as in (Kuo et al., 2022) but extended to be complementary to the target (Figure 1). Additionally, the enhancer strand has a part complementary to a biotinylated anchor. All sequences are specified in the Supplementary materials. Oligonucleotides (Integrated DNA Technologies, Eurogentec) were resuspended in Tris EDTA (TE) buffer at pH 8.0 (Sigma-Aldrich) to a stock concentration of 100 µM. AgNO_3_ (Sigma-Aldrich) was dissolved in MilliQ water to a stock concentration of 4 mM. For all further dilutions, 20 mM sodium phosphate buffer at pH 6.7−6.8, filtered using a bottle-top filter (Corning, cellulose acetate, pore size 0.22 μm), was used. Nanoclusters were made with a final concentration of templating DNA strand of 15 µM, and a ratio DNA:Ag^+^:NaBH_4_ of 1:12:24. The nanocluster-templating strand was incubated with Ag^+^ for 10 minutes in the dark, after which NaBH_4_ (Sigma-Aldrich), a reducing agent freshly dissolved in sodium phosphate buffer, was added, and the solution was shaken on a mechanical shaker for 10-20 s. The resulting solution had a pale yellow colour. The nanoclusters were left in the solution overnight in the dark at room temperature before assembling the NCBs.

NCBs were assembled at a final concentration of 1.5 µM of the templating strand with formed nanoclusters, G-rich enhancer and biotinylated anchor, while the target DNA was added at a various specified concentrations. The final sample volume was 110 µL. In negative control samples, an equal volume of the buffer was used instead of the target solution.

### Attachment of the NCBs to the NWs

After 1 hour in the dark at room temperature to allow for DNA hybridization, causing nanocluster fluorescence activation, 110 µL of NCBs were added to a flow channel (ibidi sticky-Slide VI 0.4) containing a NW platform (or planar GaP reference) functionalized with BSA-biotin and streptavidin. After 30 minutes in the dark, prior to imaging, the NCBs solution in the channel was washed away in 3 rounds with about 200 µL of 36 µM NaBH_4_ solution (NaBH_4_ was used here to maintain the solution composition), to remove unbound nanoclusters.

#### Microscopy and image analysis

We used fluorescence microscopy to measure the signal of the NCBs on NWs and on the control flat GaP substrate. Imaging was done using a Ti2-E inverted microscope (Nikon) in the flow channels with a 60X water immersion objective (Nikon) and a Sona 4.2B-11 sCMOS camera (Andor, Oxford instruments). For excitation, we used a 640 nm laser (Omicrom) at 2−6 mW of illumination power (for details, see Supporting Information). On each platform, 13 different locations (371.38 μm × 371.38 μm) were selected and imaged. The fluorescence background due to GaP autofluorescence was assessed by measuring the intensity baseline (blank) of NWs (or planar GaP reference) functionalized with biotinylated BSA and streptavidin before adding the NCBs.

For each image, 1024 × 1024 pixels (185.69 μm × 185.69 μm) sized ROIs were cropped around the center of the excitation laser, corresponding to a region with homogenous illumination. Single NW analysis was performed to quantify the fluorescence signal in each image, following a method previously developed in our laboratory (Davtyan et al., 2024). In short, all bright pixels were localized in filtered images using image dilation followed by local gradient estimation for each intensity maximum (Schnitzbauer et al., 2017). All identified maxima above local gradient threshold were fitted to Gaussian maximum likelihood estimate (MLE) model (Smith et al., 2010) to obtain subpixel position of each NW centre (Figure 2B). The average intensity *I*_*NW*_ was calculated in a 5 × 5 pixel square around the centre of each NW, after subtracting the microscope dark count. For planar reference samples, we instead measured average intensity across the platform.

## Results

We compared the signal arising from NCBs assembled in presence of the target DNA strands (Figure 1A–B) with that in absence of target (negative control). After the solution was added to the NWs and unbound DNAs were washed away, the signal was detected on the lightguiding NWs. The signal in other areas is considered as a background. It has contributions from non-bright NWs and from the areas between NWs. The first contribution is from the NWs that are not bright (Figure 2B) due to a limited quantity of assembled NCBs at a low target concentration. However, these NWs still exhibit autofluorescence. The second contribution is from the areas between NWs, and it grows with target concentration as these areas are also suitable for assembly of NCBs.

The majority of NWs show one-step photobleaching that indicates the presence of single NCBs on those NWs. The average signal from all lightguiding NWs on a platform shows an bleaching-induced exponential decay (green line in Figure 3) over the time of illumination, whereas the background signal remains essentially unchanged (in gray in Figure 3). In a region randomly chosen for photobleaching, most of NWs (58 of the 70 bright NWs detected) demonstrate a one-step bleaching (Figure S1–S2). The remaining NWs either undergo more gradual bleaching that can be attributed to multiple NCBs, or do not bleach throughout the imaging (Figures S2–S3).

**Figure 3.**
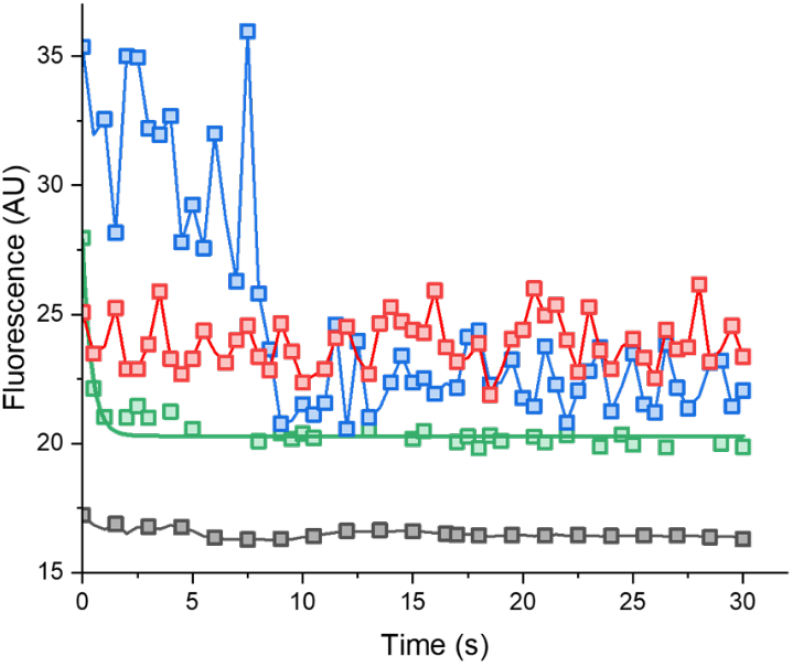
Representative fluorescence intensity curves of individual NWs (red and blue lines) on a platform where the NCBs were activated by the target DNA at 500 pM: Average signal on all lightguiding NWs (green line) and on the background (gray line). The signal on individual NWs typically bleaches in one step (blue line) or shows only marginal bleaching (red line). Photobleaching curves for other NWs are provided in Figures S1–S2.

The average fluorescence on bright NWs increases with the concentration of target DNA (Figure S4) but does not allow for reliable detection of picomolar concentrations. However, the sensitivity is improved when the number of bright NWs, another important indicator of the target presence, is taken into account. At an increasing target concentration, a larger number of NWs have bound fluorescent NCBs (Figure S5). In the approach that includes counting the NWs, the sum NW intensity *I*_*sum*_ of each image was calculated by summing the intensity of each bright NW 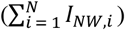 localized on a platform. The sum intensity is proportional to the number of bright NWs. Next, it was normalized by blank intensity, that is, the sum intensity on NWs on the same area of that platform before adding the NCBs. That way, we accounted also for the autofluorescence signal from the blank platforms. We then used the sum intensities to calculate the fluorescence relative to the signal from a platform where no target was added (negative control). The absence of target means that NBCs should not assemble, hence nanoclusters could only bind to the platform in a non-specific manner and without proper activation of their fluorescence by G-rich enhancer strands.

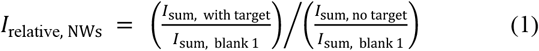

The relative signal is thus corrected for both autofluorescence of the platform and baseline fluorescence in absence of target. To compare this to detection of the same target by NCBs not on NWs but on planar GaP, we calculated *I* relative for the planar platforms where we measured the average intensity across the platform.

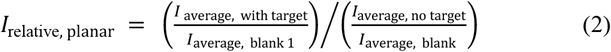

The results (Figure 4) show detection already at picomolar concentrations of the target DNA, whereas on planar GaP we do not achieve detection even at 0.1 µM of target. Thus, the NW platforms lower the limit of detection by at least five orders of magnitude.

**Figure 4.**
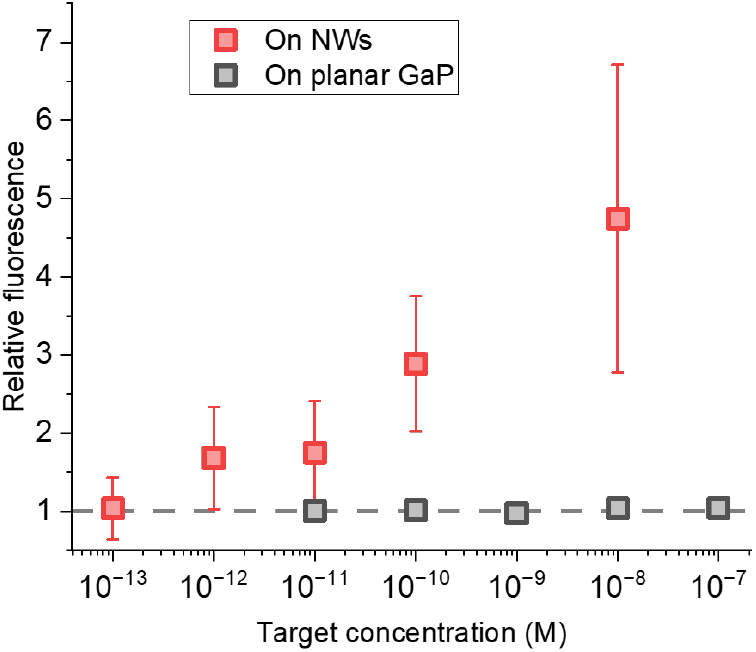
Relative fluorescence on NWs and planar GaP at a varied concentration of target. The relative signal was calculated according to the Equations (1) for the NWs and (2) for planar GaP. Uncertainties were calculated as standard deviation between locations on the platforms.

The sensitivity indicated by the intensity contrast between the NCBs assembled in presence of target and the negative control is thus higher when estimated from the sum intensity on NWs than from average intensity. A more prominent contrast between the sum intensities is due to a significantly larger number of bright NWs (Figure S5) detected on the platform where the target was present.

## Discussion

The method we present here is simple to implement: the end user only needs to add a sample to a solution of NC-templating, enhancer and stabilizer strands, which can be supplied pre-mixed. The resulting mixture is then added to a flow channel with NWs, and fluorescence is detected after a single washing step. No costly reagents or target amplification are required, while short DNA oligos are inexpensive.

Table 1 summarizes key differences between this sensor and PCR. PCR detection of short targets (less than 50 bases) relies on even shorter primers to start elongation. This generally decreases specificity of amplification (Grunenwald, 2003). Although our method also requires hybridization of short parts of the target with the NCB strands, the detection does not involve amplification and is expected to only take place when the entire target is complementary to the respective strands (Table 1). Unlike in previous studies (Kuo et al., 2022; Obliosca et al., 2014; Zou et al., 2019), we assemble NCBs at room temperature which simplifies the process.

**Table 1.** Requirements and limitations of the NCB-NW sensor as compared to PCR. The information about PCR from (Bruce et al., 2020; Harikai et al., 2022; Iwanaga, 2022; Madadelahi et al., 2024; Nakano et al., 2017) is used.

|  | Sensitivity | Equipment and materials | Time | Targets |
| --- | --- | --- | --- | --- |
| NCB-NW sensor | Low picomolar | Fluorescence microscope, NWs, three oligos including a nanocluster-templating, silver salt, reducing agent | DNA hybridization + 30 minutes for binding to NWs | Short single stranded DNA (20-50 bases) at room temperature, potentially – longer targets with annealing |
| PCR | Femtomolar to low picomolar | PCR machine, primers, DNA polymerase, dNTPs, reverse transcriptase for RT-PCR | Minutes (fast) to hours (slow) | Typically longer than 40-50 bases as shorter targets require careful optimization or advanced methods. Samples must be free from PCR inhibitors. |

We leverage a key advantage of NW platforms for biosensing, namely that each of multiple simultaneously imaged NWs can enhance and guide the signal of surface-bound fluorophores. This signal is multiplied by a large number of the NWs, making platforms with high NW density preferred.

The photobleaching traces (Figure 3 and S3) indicate that our method has single-nanocluster sensitivity. This is supported by the fact that the platforms remain unsaturated by the NCBs (Figure 2), that is, only a fraction of all NWs on a platform get an assembled NCB on them at the concentrations used here. This also suggests a broad dynamic range of detection (Davtyan et al., 2024). In terms of sensitivity, our approach surpasses the electrochemical (Zhang et al., 2010) and fluorimetric (Zou et al., 2019) detection previously used for a similar target DNA.

In our sensor, only the non-fluorescent part of a beacon, that is, a G-rich enhancer strand is immobilized on the surface so that silver nanoclusters do not remain in proximity of NWs unless the target DNA is present. This reduces the signal in absence of a target.

## Author contributions

I.N.U. wrote the manuscript with input from co-authors. I.N.U. and H.L. conceptualized the study and designed experiments with advice from C.N.P.. I.N.U. and R.D. performed the experiments and analyzed the data.

## ACKNOWLEDGMENT

For financial support, we thank NanoLund (Junior Scientist Ideas Award grant to I.N.U.), the Swedish Research Council (Project 2020-04226), and the European Union’s Horizon 2020 research and innovation programme under the Marie Skłodowska-Curie grant agreement No 945378 (to R.D.) NWs were grown and characterized by Aligned Bio AB in Lund Nano Lab (LNL). We are also grateful to Prof. Fredrik Höök, Prof. Ken’ya Furuta, Prof. Thoas Fioretos, Assoc. Prof. Nicklas Anttu and Dr. Nils Gustafsson for useful discussions.

