## Supplementary for "Nanowire-based biosensor for short DNA using fluorescent silver nanoclusters"

### Supplementary data

### Illumination conditions

The laser illumination power was measured using the power meter Thorlabs PM100D placed on top of the used 60X water immersion objective (Nikon) without adding water, in the light path within the microscope. It was measured at 15−40% of the maximum power at 16 mW, that is, 2−6 mW.

### Assembling the nanocluster beacons

The oligonucleotide sequences were as follows (5’ to 3’, complementary parts have the same color):

Target HIV-1: GCTATACATTCTTACT ATTTTATTTAATCCCAG

Biotinylated G-rich enhancer: biotin-CGTGTAGGTCATAGAT CTGGGATTAAATAAAAT TCCATTGGTGGGGTGGGG

NC-templating strand: CCCCCTTAATCCCCC AGTAAGAATGTATAGC

Stabilizer strand: ATCTATGACCTACACG

The enhancer and templating sequences are identical to those in (Kuo et al., 2022), the HIV-1 sequence is identical to that in (Zhang et al., 2010).

#### Photobleaching on individual NWs

We monitored the intensity of lightguiding NWs and background upon continuous illumination.

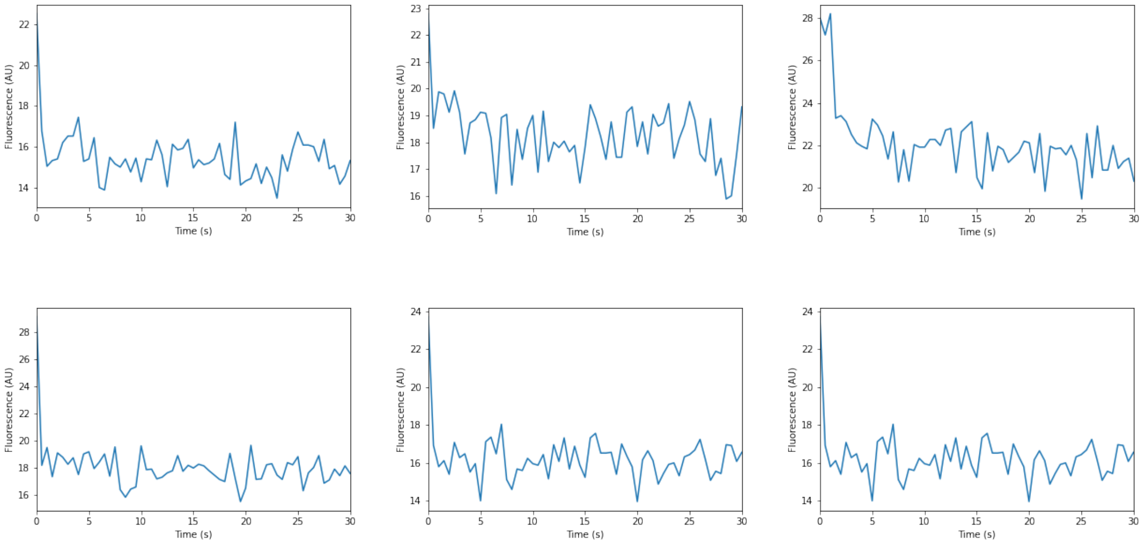


Figure 1. Representative fluorescence intensity curves from exemplary NWs on a platform, which show one-step photobleaching.

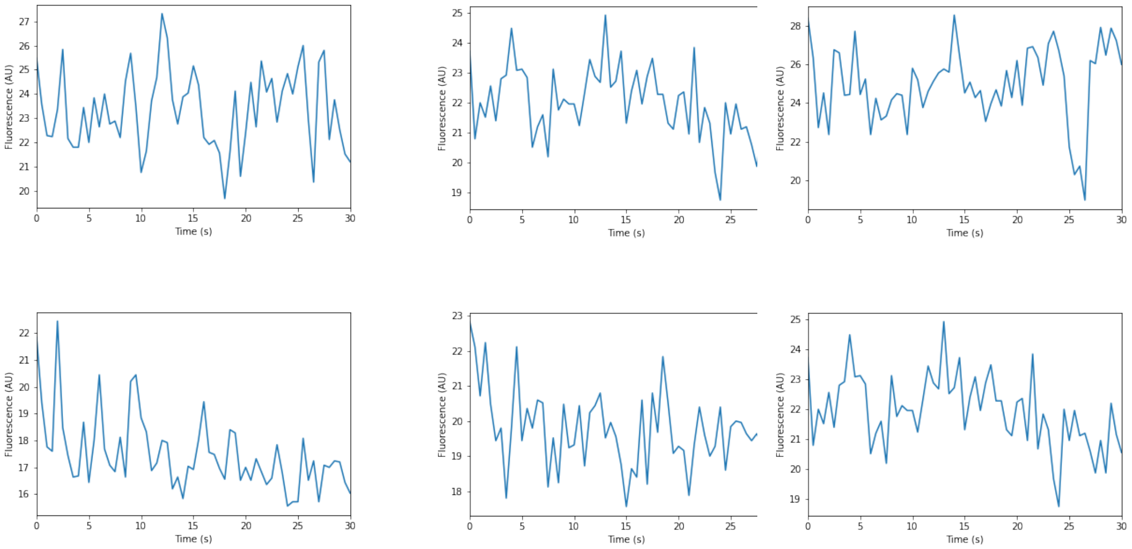


Figure 2. Representative fluorescence intensity curves from NWs on a platform, which show gradual photobleaching or no significant intensity changes.

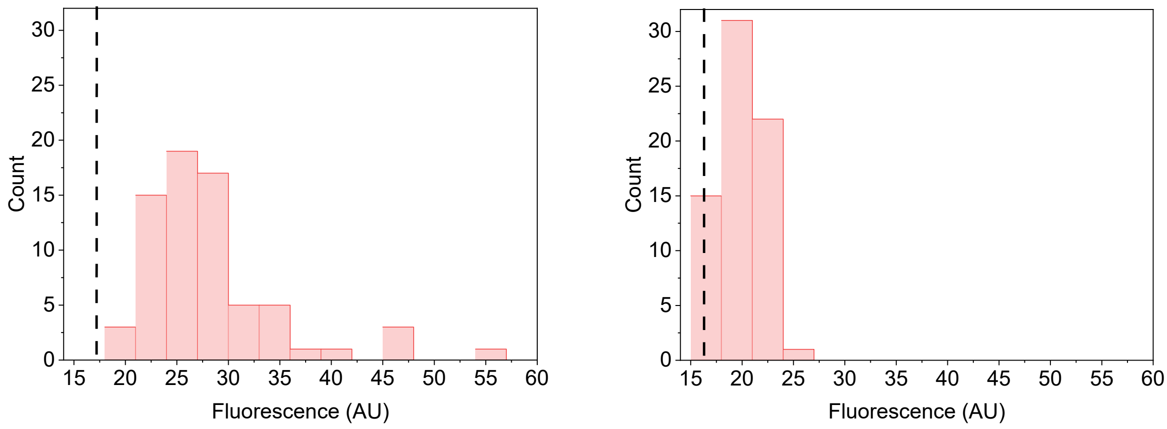


Figure 3. Fluorescence intensities (A) in the beginning and (B) after 30 s of illumination. Black dashed line shows the background signal for the same time points.

### Average intensity on lightguiding NWs and background

The average intensities were calculated for all NWs detected as bright in fluorescence images. The background average intensity was calculated for all other regions within the field of view.

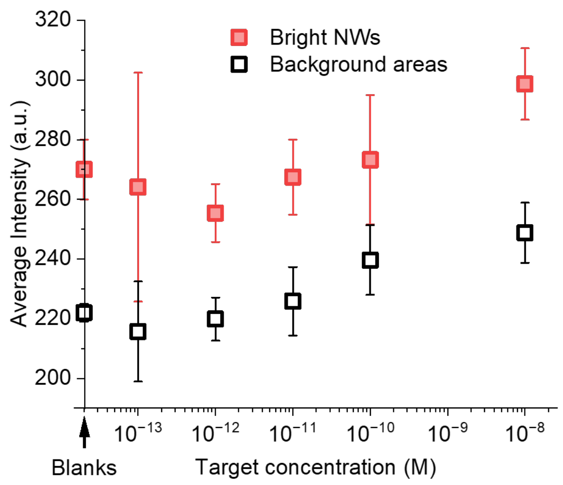


Figure 4. Average fluorescence on platforms where NCBs were added at a varied concentration of the DNA target, and on blank platforms where no NCBs were added (symbols on the y-axis). The intensities were measured on bright NWs and other areas (background). Uncertainties were calculated as standard deviation between the NWs.

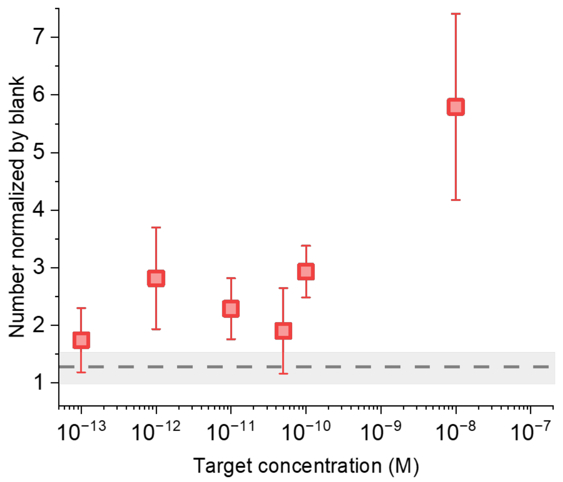


Figure 5. Number of bright NWs with activated NCBs. The dashed line and respective uncertainty indicate the normalized number of bright NWs in the platforms where no target was added. Uncertainties were calculated as standard deviation between locations on the platforms.
